# cGAS-activating lupus autoantibody for cancer immunotherapy

**DOI:** 10.1101/2023.02.11.527649

**Authors:** Xiaoyong Chen, Xiangjun Tang, Benedette J. Cuffari, Caroline Tang, Xingchun Gao, Philip W. Noble, Melissa R. Young, Olivia M. Turk, Anupama Shirali, Joseph Gera, Robert N. Nishimura, Jiangbing Zhou, James E. Hansen

## Abstract

Cytoplasmic DNA triggers a cGAS-mediated signaling cascade that promotes an innate immune response and is potentially actionable in cancer immunotherapy. Here we show that a cytoplasmic-localizing lupus anti-DNA autoantibody activates cGAS and facilitates an immune-mediated prolongation of survival in orthotopic models of glioblastoma (GBM). Mechanistically, cellular penetration and blood-brain barrier crossing by the anti-DNA autoantibody is linked to nucleoside transport. Pulldown, knockdown, signaling, and cytotoxicity assays demonstrate autoantibody association with and activation of cGAS. In orthotopic GBM models, the autoantibody localizes to brain tumor, increases tumor CD8+ T cell content, and prolongs survival in immunocompetent but not immunodeficient mice. This work introduces the new concept of a cGAS-activating anti-DNA autoantibody, which impacts theories on mechanisms of autoimmunity and has translational applications in cancer immunotherapy.

## Introduction

Anti-DNA antibodies in the autoimmune disease systemic lupus erythematosus (SLE) are protective against some malignancies (1, 2). Cell-penetrating variants of anti-DNA autoantibodies have emerged as an unexpected source of biologics capable of engaging hard to reach intracellular targets and exerting synthetically lethal effects on vulnerable cancer cells. Nuclear-localizing and DNA-damaging autoantibodies, referred to as Deoxymabs, are in clinical trial planning for use against DNA repair-deficient malignancies (2-5). In contrast to Deoxymabs that enter nuclei and act on nuclear DNA, other anti-DNA autoantibodies localize into the cytoplasm (6). Presence of cytoplasmic DNA triggers an inflammatory cascade that contributes to mechanisms of immunity (7), and in the present work we find that a cytoplasmic-localizing anti-DNA autoantibody augments this response and offers a new strategy in cancer immunotherapy.

Cyclic GMP-AMP synthase (cGAS) is a cytoplasmic DNA sensor that plays a central role in innate immunity by producing cyclic GMP-AMP (cGAMP) to activate stimulator of interferon genes (STING) signaling and a type I interferon response (7). The cGAS-STING pathway is potentially actionable in approaches to cancer immunotherapy (8, 9), and we hypothesize that cytoplasmic penetrating anti-DNA autoantibodies can impact cGAS function. Some anti-DNA autoantibodies preferentially localize to microenvironments enriched in DNA released by malignant and necrotic tissue, which raises the possibility that they may be used to selectively target tumors (5, 10-12). This strategy is exemplified in Deoxymab-1 (DX1), which uses a nucleoside transport-dependent mechanism of cell membrane transit, including crossing of the blood-brain barrier (BBB). As a derivative of an anti-DNA autoantibody, DX1 accumulates at necrotic tumor environments enriched in extracellular DNA and nucleosides and is subsequently driven into live tumor cells through nucleoside transporters salvaging local DNA. DX1 is a compelling agent for strategies targeting intranuclear processes in central nervous system (CNS) malignancies and is in preparation for clinical trials against glioblastoma (GBM) (5). In the present work we sought to find a cytoplasmic localizing anti-DNA autoantibody that uses a nucleoside transport-dependent mechanism of cellular penetration and BBB crossing, like DX1, and to evaluate its potential to activate cGAS in an approach to brain tumor immunotherapy.

Anti-DNA autoantibodies are widely diverse with varying nucleic acid binding specificities, with some recognizing multiple conformations and sequences of DNA and others seeking epitopes on specific nucleosides (13-15). Guanosine (GUO) is the most immunogenic nucleoside, and of all anti-DNA autoantibodies produced in lupus the anti-GUO titers correlate most accurately with disease severity (16, 17). Functionally, anti-GUO autoantibodies share a common binding motif with guanine nucleotide-binding proteins and may impact G-protein activity and associated signaling pathways (6, 18, 19). Previous work identified a 4H2 anti-DNA autoantibody that binds an epitope on GUO, penetrates live cells, and localizes into cytoplasm. Moreover, nucleoside transport was suggested to be involved in its mechanism of uptake (6, 18, 20). We selected 4H2 as a prototype cytoplasmic-localizing anti-DNA autoantibody and examined its impact on cGAS function, patterns of transport into cells and across the BBB, and potential to enhance tumor immune response in orthotopic models of GBM.

## Results

### 4H2 penetrates glioma cells, localizes to cytoplasm, and associates with cGAS

4H2 purified from hybridoma supernatant migrated as expected on SDS-PAGE (**Supp. Fig. 1a**), and antibody penetration into U87 and PPQ glioma cells was evaluated. U87 is a commonly used human glioma cell line (21), and PPQ cells are murine glioma cells with deletions in *TRP53, Pten*, and *Qk* and are syngeneic with and form highly invasive GBM tumors in the brains of C57/BL6 mice (22). U87 and PPQ cells were treated with IgG control, DX1, or 4H2 and probed to detect penetration by immunofluorescence. No significant cellular uptake of IgG control was observed, while DX1 penetrated nuclei as expected. In contrast, 4H2 localized into the cytoplasm of the glioma cells (**Fig. 1**).

**Figure 1.**
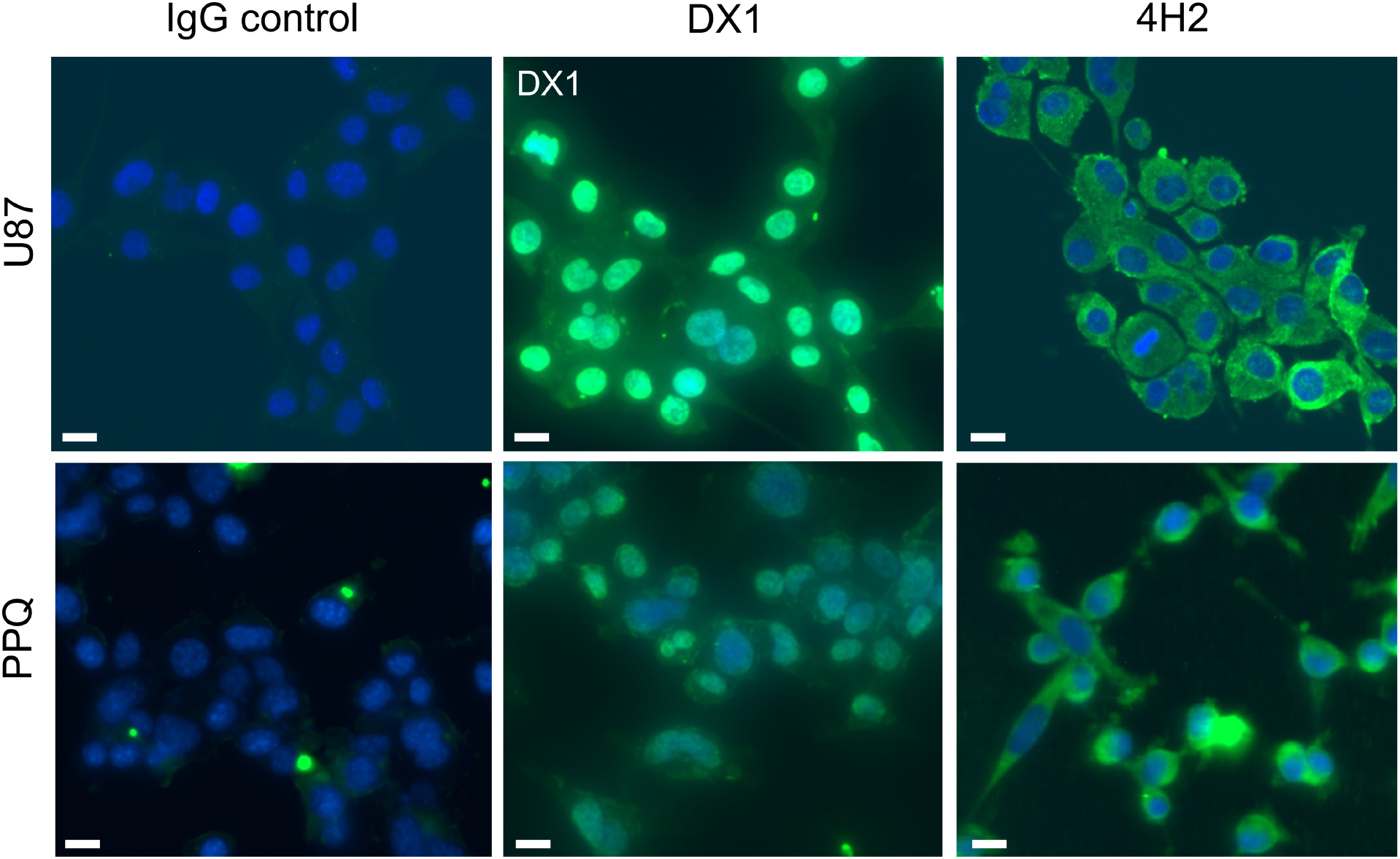
4H2 localizes into the cytoplasm of glioma cells. Human U87 and murine PPQ glioma cells treated with IgG control, DX1, or 4H2 were immunostained to detect antibody penetration and counterstained with DAPI. No significant uptake of IgG control was observed in either cell line. DX1 and 4H2 penetrated and localized into the nuclei and cytoplasm of cells, respectively. Representative merged images are shown. Bar = 10 µm.

Both cGAS and 4H2 bind DNA, and we hypothesized that 4H2 would associate with cGAS either through direct binding or through simultaneous binding to an intermediary nucleic acid. Purified recombinant cGAS was incubated with IgG control or 4H2 ± nucleic acid, and antibodies and their bound protein were isolated using protein G. 4H2 showed greater association with purified cGAS compared to protein G/IgG control, and addition of nucleic acid reduced the 4H2-cGAS interaction but not the nonspecific protein G/IgG control-cGAS association (**Fig 2a**). These findings suggest a 4H2-cGAS interface that is competitively inhibited by nucleic acid. Equivalent IgG control and 4H2 content in the pulldown samples was confirmed by anti-IgG western blot (**Fig 2b**). To probe for an interaction between 4H2 and cGAS in cells, the protein G-based antibody pulldown assay was performed on lysates of PPQ cells treated with IgG control or 4H2. Anti-cGAS western blot showed greater cGAS pulldown in 4H2-treated cells compared to IgG control (**Fig. 2c**). Ras was used as a negative control G-protein, and no binding by IgG control or 4H2 to Ras was detected (**Fig. 2d**). This finding correlates with the results using purified cGAS and demonstrates an association between 4H2 and cGAS in cells.

**Figure 2.**
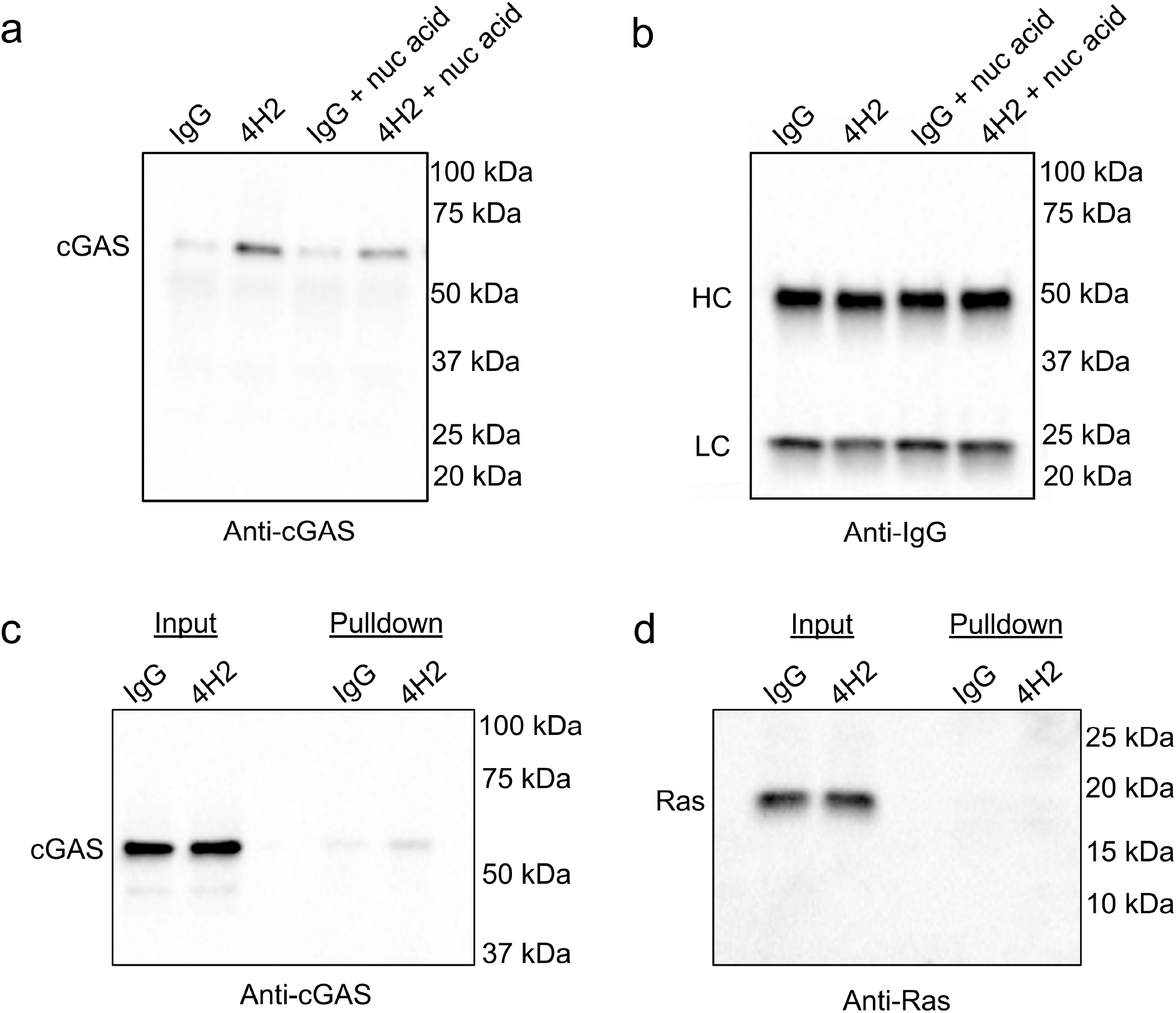
4H2 interacts with cGAS. (**a, b**) 4H2 associates with purified cGAS. Purified recombinant cGAS ± nucleic acid was incubated with IgG control or 4H2. Antibodies and bound protein were then isolated using protein G beads. 4H2 showed increased association with cGAS compared to protein G/IgG control. Presence of nucleic acid (nuc acid) reduced the 4H2-cGAS interaction, consistent with competitive inhibition, but did not impact the nonspecific association of protein G/IgG control with cGAS (**a**). Equivalent IgG control and 4H2 content in the protein G pulldown samples was confirmed by anti-IgG western blot (**b**). (**c, d**) 4H2 associates with cGAS in PPQ cells. Antibodies and associated bound proteins were isolated from lysates of IgG control or 4H2-treated PPQ cells using protein G beads. Western blots of input lysate and protein G pulldown showed greater cGAS pulldown in 4H2-treated cells compared to IgG control (**c**), while binding to negative control Ras was not detected (**d**). All blots are representative of three independent experiments.

### 4H2 enhances cGAS activity

Activated cGAS catalyzes the formation of cGAMP (7). cGAMP production by cGAS was increased 83%±18 in the presence of 4H2 compared to IgG control *in vitro* (P<0.05) (**Fig. 3a**), which demonstrates 4H2 enhancement of cGAS activity. NF-κB nuclear translocation was subsequently used as a marker of cGAS activation to examine the impact of 4H2 on cGAS function in cells. Western blots of cytoplasmic and nuclear contents of PPQ cells revealed an increase in NF-κB nuclear content by a factor of 2.2±0.2 in 4H2 compared to IgG control-treated cells (P<0.05) (**Fig. 3b, c**), consistent with 4H2-mediated cGAS activation.

**Figure 3.**
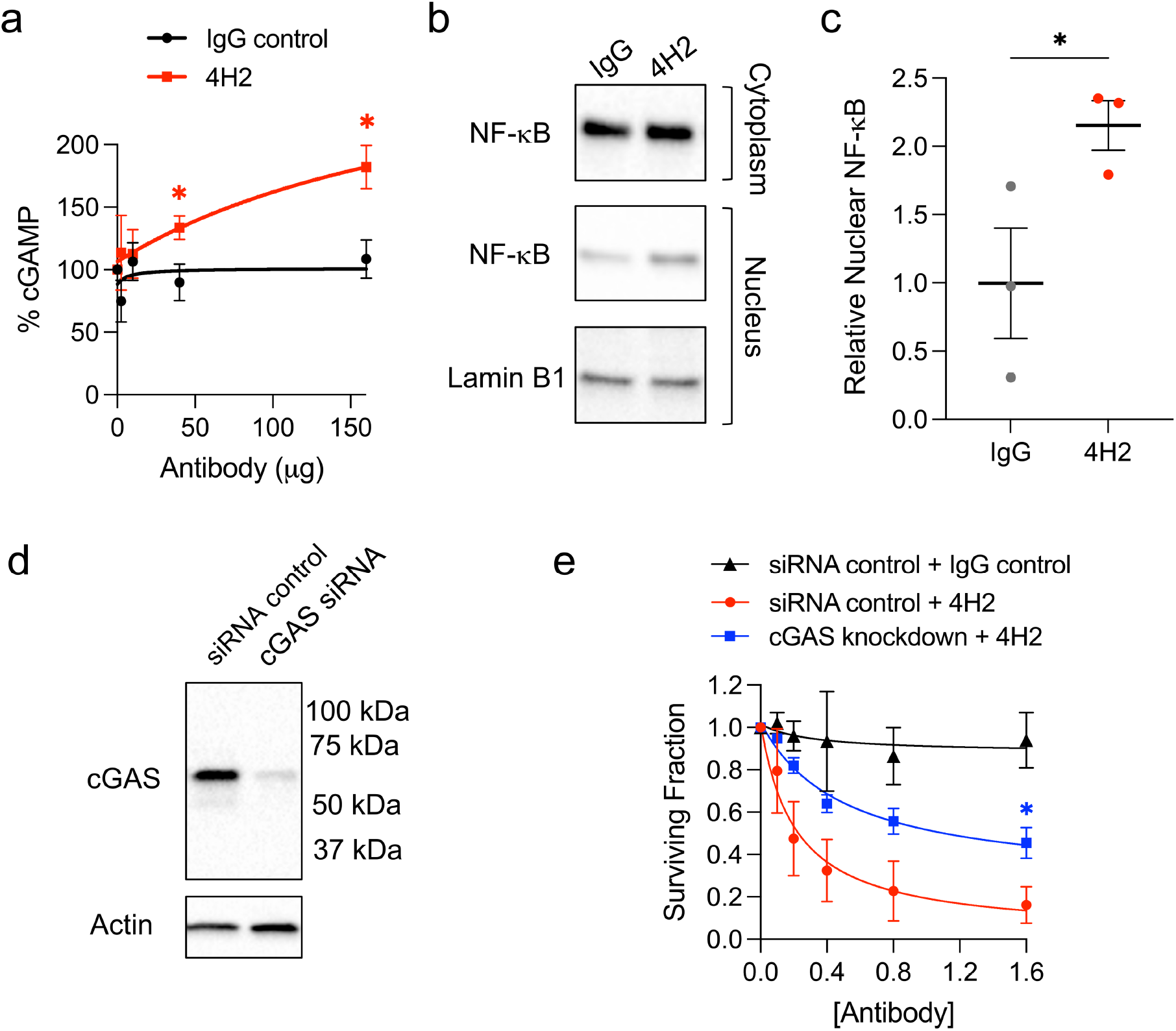
4H2 enhances cGAS activity. (**a**) 4H2 causes a dose-dependent increase in cGAS activity. The ratio of cGAMP production by cGAS in the presence of 4H2 or IgG control was used to determine the percentage increase in activity (*P<0.05). (**b, c**) 4H2 induces nuclear translocation of NF-κB. Cytoplasmic and nuclear contents of PPQ cells treated with IgG control or 4H2 were analyzed by western blot probed for NF-κB, and Lamin B1 for loading control. Representative blots are shown in (**b**), and relative NF-κB nuclear content quantified by ImageJ in (**c**). 4H2 increased relative nuclear NF-κB by a factor of 2.2±0.2 (*P<0.05). (**d, e**) 4H2 mediates cGAS-dependent cytotoxicity. PPQ cells transfected with control or cGAS siRNA were treated with IgG control or 4H2. cGAS western blot confirmed successful knockdown (**d**), and colony formation assay demonstrated cGAS-dependent toxicity of 4H2 (**e**). (*P<0.05).

Previous work has shown that activation of cGAS promotes cellular senescence and apoptosis (23, 24). To further probe the interaction between 4H2 and cGAS, control or cGAS-knockdown PPQ cells were treated with IgG control or 4H2 (0-1.6 µM) and surviving fractions determined by colony formation assay. IgG control was not significantly toxic to the cells, with surviving fraction 0.94±0.13 (ns) after treatment at the highest dose tested of 1.6 µM. In contrast, 4H2 was highly toxic to the cells, and cGAS knockdown significantly reduced cellular sensitivity to 4H2. Knockdown was confirmed by western blot (**Fig. 3d**) and surviving fractions were 0.16±0.09 and 0.46±0.07 in control or cGAS-knockdown PPQ cells treated with 1.6 µM 4H2, respectively (P<0.05) (**Fig. 3e**). Taken together, these findings demonstrate an enhancing interaction between 4H2 and cGAS and a cGAS-dependent toxicity of 4H2 to glioma cells.

### 4H2 exhibits a nucleoside-dependent mechanism of transport

As previously mentioned, the DNA-binding and nucleoside transport-dependent mechanism of membrane transit by DX1 facilitates its crossing of the BBB and its preferential localization into tumors (5). Previous work by others suggested a similar role for nucleoside transporters in 4H2 uptake (20). While DX1 and 4H2 are both cell-penetrating anti-DNA autoantibodies, they are distinct in their patterns of intracellular localization and DNA binding. DX1 has an affinity for a range of DNA substrates and penetrates nuclei, whereas 4H2 specifically binds an epitope on GUO and localizes into cytoplasm. To test the role of nucleosides in 4H2 uptake PPQ cells treated with 4H2 ± adenosine (ADE) or GUO, or positive control DX1 ± adenosine (ADE), were immunostained to detect antibody penetration. Efficiency of membrane crossing was determined by ImageJ quantification of fluorescence signal. DX1 responded to ADE as expected, with nuclear penetration enhanced by a factor of 1.70±0.01 (P<0.0001) (**Fig. 4a**). ADE did not enhance 4H2 uptake, with cytoplasmic signal 0.97±0.16 in the presence of ADE relative to its absence (ns) (**Fig. 4b**). In contrast, GUO increased 4H2 penetration by a factor of 3.2±0.3 (P<0.0001) (**Fig. 4c**). These findings demonstrate a GUO-responsive mechanism of cellular penetration by 4H2 that correlates with the known affinity of 4H2 for GUO and implicates nucleoside transport in 4H2 uptake.

**Figure 4.**
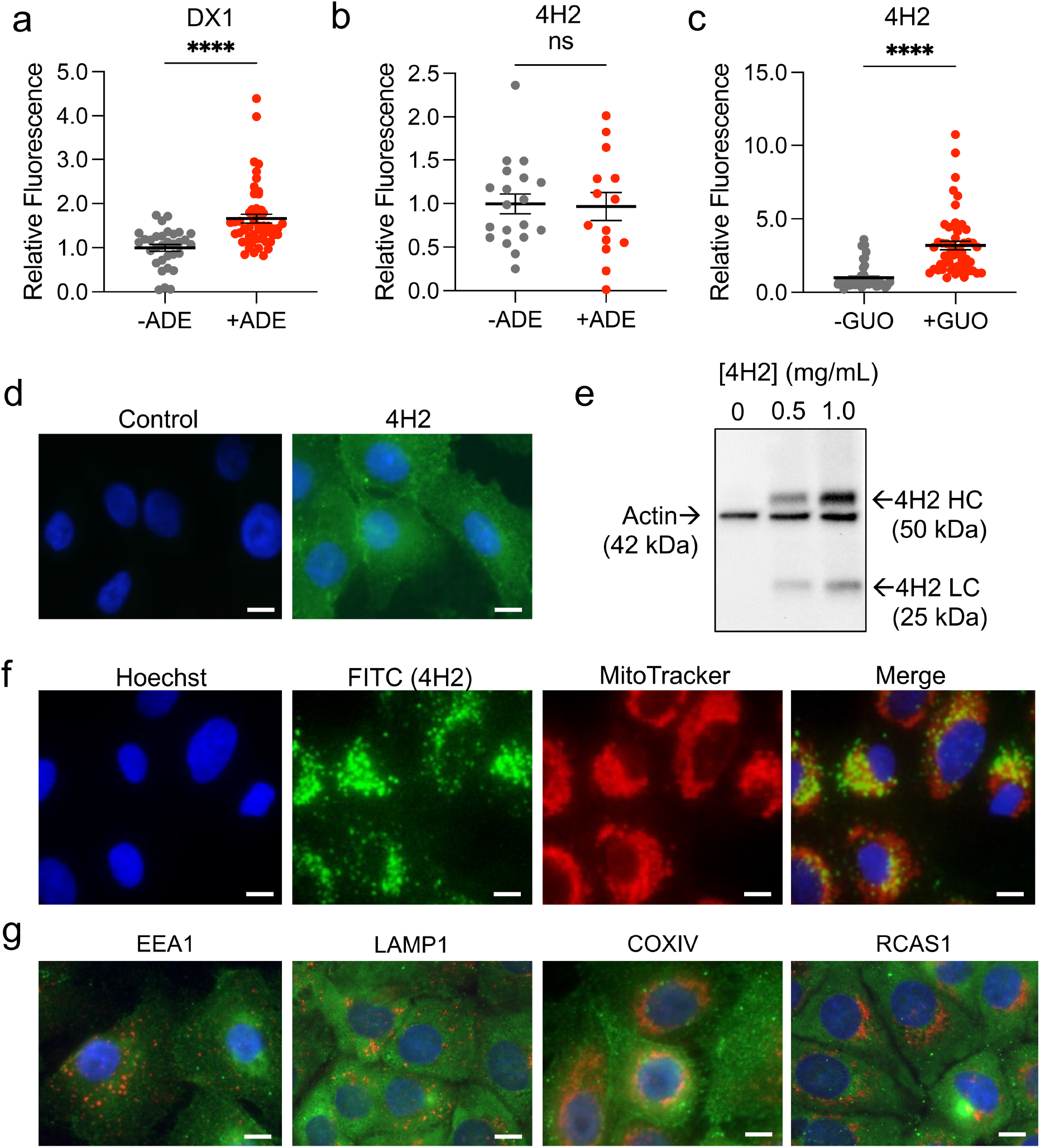
4H2 penetrates cells through a GUO-responsive mechanism and avoids endosomes and lysosomes. (**a-c**) The effect of supplemental nucleoside on cellular penetration by DX1 and 4H2 into PPQ cells was determined by ImageJ quantification of fluorescence signal. ADE supplementation increased the efficiency of cellular penetration by DX1 by a factor of 1.70±0.01 (****P<0.0001) (**a**) but did not enhance 4H2 penetration (**b**). In contrast, GUO supplementation increased 4H2 cellular penetration by factor of 3.20±0.30 (****P<0.0001) (**c**), correlating with the known affinity of 4H2 for GUO. (**d**) 4H2 penetrates lung adenocarcinoma cells and localizes into cytoplasm. Representative images of control and 4H2-treated Cal12T cells immunostained for antibody penetration and counterstained with DAPI are shown. (**e**) Lysates of Cal12T cells treated with 0-1 mg/mL 4H2 for 24 hours were analyzed by western blot probed with primary actin antibody for loading control and an anti-mouse secondary antibody to detect the actin primary and 4H2 (both murine). 4H2 HC and LC ran at their expected MWs, showing the antibody is not significantly degraded 24 hours after cellular penetration. (**f**) Visualization of live A549 lung adenocarcinoma cells treated with FITC-labeled 4H2 and MitoTracker Red FM and Hoechst counterstains show 4H2 penetration into cytoplasm. Images of Cal12T cells from this experiment are shown in **Supp. Fig. 1b**. (**g**) Representative images of Cal12T cells treated with 4H2 immunostained for 4H2 (green) and markers of specific organelles (red), including EEA1 for endosomes, LAMP1 for lysosomes, COX IV for mitochondria, and RCAS1 for Golgi apparatus), and DAPI nuclear counterstained (blue). 4H2 signal did not specifically co-localize with any of the organelles, including endosomes and lysosomes. Error bars = SEM. Scale bars = 10 µm.

### 4H2 avoids endosomes and lysosomes

Mechanisms of cellular uptake by 4H2 were further probed in lung adenocarcinoma cells (A549 and Cal12T) with cytoplasmic compartments that are more readily visualized under microscopy compared to glioma cells. 4H2 penetrated lung cancer cells and localized to cytoplasm (**Fig. 4d**), and analysis of cell lysates twenty-four hours after treatment with 4H2 by western blot probed with anti-mouse IgG secondary revealed 4H2 heavy chain (HC) and light chain (LC) at their expected MW. This demonstrated retention of intact antibody chains twenty-four hours after penetrating cells (**Fig. 4e**). Any contribution of fixation artifact was ruled out by live cell imaging of lung cancer cells treated with FITC-labeled 4H2 and counterstained with MitoTracker Red FM and Hoechst 33342 (ThermoFisher Scientific, Waltham, MA) (**Fig. 4f and Supp. Fig. 1b**). Overlay of FITC, Hoechst, and MitoTracker images showed an absence of 4H2 FITC signal in the nuclei consistent with previous reports (6).

The stability of 4H2 after penetrating cells suggested an avoidance of endosomes and lysosomes (**Fig. 4e**). Fluorescence co-localization studies probed the intracellular location of 4H2 after its uptake into cells. Cells treated with 4H2 were immunostained with Alexa Fluor 488-conjugated anti-mouse IgG antibody to detect 4H2 and rabbit primary antibodies to detect markers of early endosomes (EEA1), lysosomes (LAMP1), Golgi (RASC1), endoplasmic reticulum (PDI), and mitochondria (COX IV), followed by Alexa Fluor 555-conjugated anti-rabbit IgG antibody and DAPI nuclear counterstain (Cell Signaling Technology, Danvers, MA). 4H2 did not co-localize with endosomes or lysosomes, or any of the other organelles tested (**Fig. 4g**).

### 4H2 penetrates cells and crosses a transwell model of the BBB through a nucleoside transporter-dependent mechanism

The lack of early degradation and the avoidance of endosomes/lysosomes by 4H2 observed here suggests a non-endocytic mechanism of cellular penetration. Further, the enhancing effect of GUO on 4H2 penetration described above implicates a nucleoside transport-based mechanism, which is consistent with a previous abstract report (20). We further tested the dependence of 4H2 transport on nucleoside transport by treating lung cancer cells with the nucleoside transport inhibitor dipyridamole (DP) and examining its effect on subsequent efficiency of cellular penetration by 4H2. DP significantly inhibited 4H2 uptake, with relative signal as quantified by ImageJ reduced to 0.29±0.03 (P<0.0001) compared to 4H2 penetration in the absence of DP (**Fig. 5a**).

**Figure 5.**
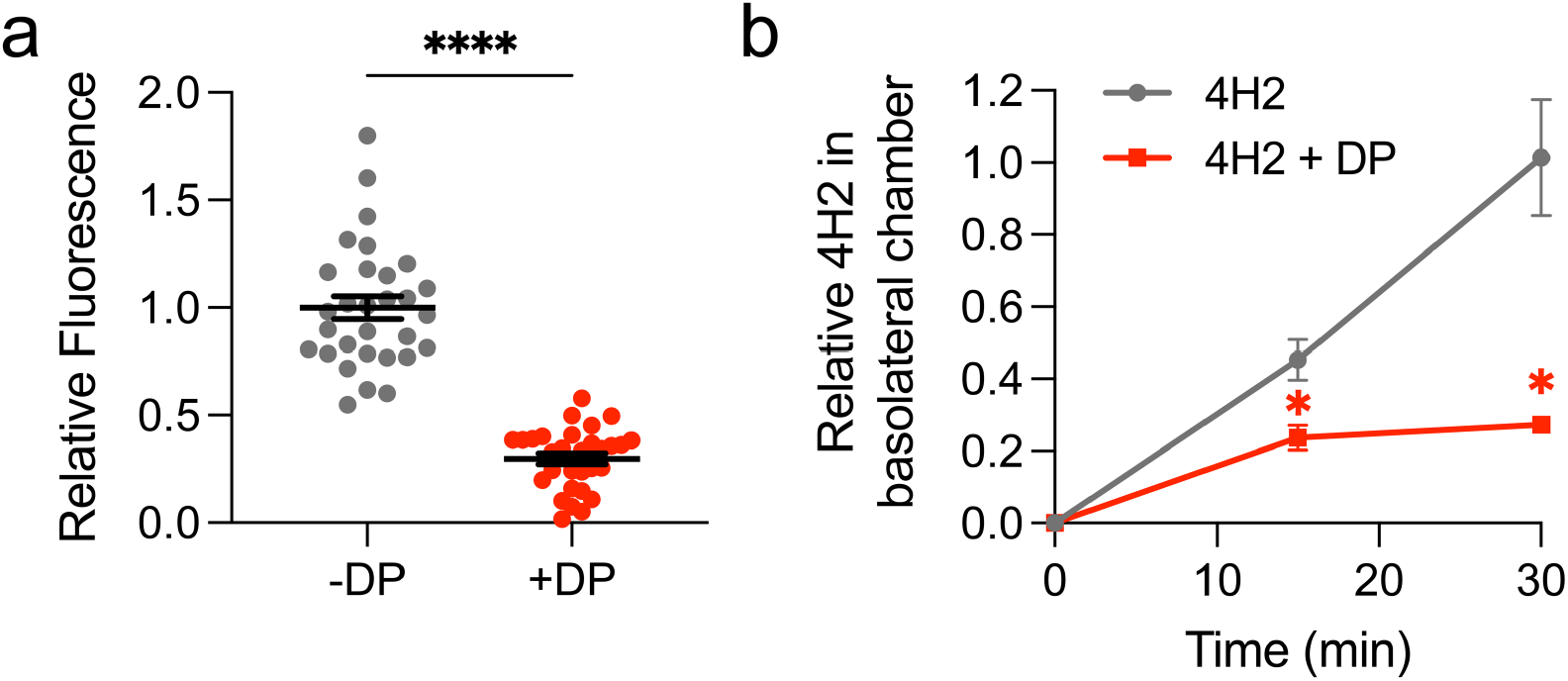
Nucleoside transporter-dependent membrane transit by 4H2. (**a**) Inhibition of nucleoside transport blocks 4H2 cell penetration. 4H2 penetration into Cal12T cells by 4H2 ± 50 µM DP was evaluated by ImageJ quantification of fluorescence signal. DP reduced 4H2 penetration to 0.29±0.03 (****P<0.0001) relative to 4H2 uptake in the absence of 4H2. (**b**) A transwell model of the BBB using hCMEC/D3 BECs and NHAs was used to evaluate 4H2 transit across the barrier from apical to basolateral chambers. 4H2 crossed the barrier, and transport was suppressed by the nucleoside transport inhibitor DP. Relative basolateral content of 4H2 at thirty minutes after application into the apical chamber was reduced to 0.273±0.003 (*P<0.05) in the presence of 50 µM DP.

Nucleoside transport facilitates BBB crossing and brain tumor localization by DX1, and DP has previously been shown to block this process (5). Given similarities in mechanism of cellular penetration we hypothesized that 4H2 would also cross the BBB and penetrate brain tumors. 4H2 transit across an hCMEC/D3 human brain endothelial cell-based transwell model of the BBB was evaluated using the same protocol previously used for DX1 (5). 4H2 successfully crossed the transwell BBB model to move from apical to basolateral chambers. In addition, treatment of the BBB with the nucleoside transport inhibitor DP prior to addition of 4H2 to apical chambers reduced 4H2 crossing into the basolateral chamber, indicating nucleoside transport-dependent transit (**Fig. 5b**). Taken together, these findings indicate that 4H2, like DX1, uses a nucleoside transporter-dependent mechanism of transport across cell membranes.

### 4H2 localizes into orthotopic GBM and prolongs survival in syngeneic GBM models

DX1 localizes into orthotopic GBM through a nucleoside transport-dependent mechanism (5). Based on the findings above, we expected 4H2 would similarly localize into orthotopic GBM. C57/BL6 mice with syngeneic orthotopic PPQ GBM brain tumors confirmed by IVIS were treated with a single dose of IgG control or 4H2 at 40 mg/kg (dose extrapolated from previous work with DX1) by tail vein injection. Tumors and normal tissues were taken after twenty-four hours, and presence of antibody evaluated by immunohistochemistry (IHC). Significant antibody staining was detected in the cytoplasm of GBM tumor cells after treatment with 4H2, but not IgG control (**Fig. 6a**). No antibody stain was detected in normal brain tissue remote from the tumors in mice treated with IgG control or 4H2 (**Fig. 6b**). In normal tissues, 4H2 showed an increased localization to kidney compared to IgG control, but otherwise similar distribution of antibodies was seen in skeletal muscle (**Fig. 6c, d**). This pattern of localization by 4H2 to brain tumor but not normal brain parallels the previously reported activity of DX1 (5) and is consistent with a DNA/nucleoside-facilitated mechanism of uptake.

**Figure 6.**
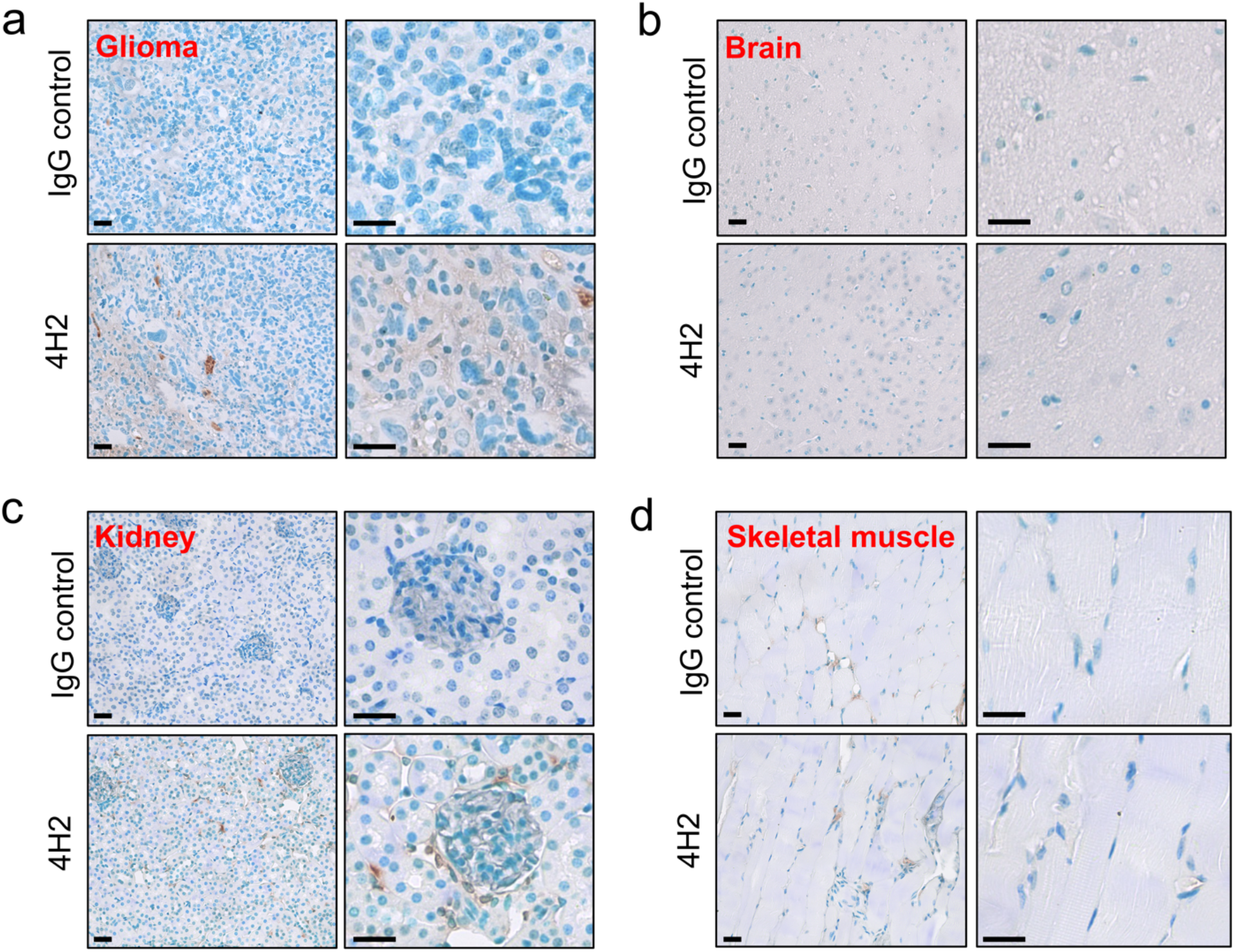
4H2 localizes to orthotopic GBM. (**a-d**) Tissue localization by systemically administered antibodies in C57/BL6 mice bearing PPQ orthotopic GBM tumors was evaluated by IHC twenty-four hours after treatment with IgG control or 4H2. Stain consistent with cytoplasmic localization of antibody was observed in the GBM tissue (**a**) but not normal brain remote from tumor after treatment with 4H2 (**b**), while minimal stain was detected in tumors or normal brain after treatment with IgG control (**a, b**). 4H2 treatment was associated with a small increase in antibody detected in kidneys compared to IgG control (**c**), while no difference was detected between 4H2 and IgG control in skeletal muscle (**d**). This pattern of 4H2 localization to tumor but not normal brain is consistent with DNA/nucleoside-facilitated uptake.

Mice bearing orthotopic PPQ GBM tumors were randomized to treatment with IgG control (N=4) or 4H2 (N=5) at 40 mg/kg by tail vein injection weekly for three weeks and monitored for toxicity and survival. No adverse effects were observed. 4H2 increased median survival by 66% compared to mice treated with IgG control (P<0.01, log-rank test). Survival to study completion was 40% in the group treated with 4H2 and 0% in the IgG control group (**Fig. 7a, Supp. Table 1**). Mechanistically, we hypothesized the prolonged survival mediated by 4H2 to be a consequence of cGAS-mediated senescence/toxicity to GBM tumor cells, cGAS-mediated enhancement of immune response, or a combination thereof. Tumors from mice meeting criteria for euthanasia (N=4 and 3 for IgG control and 4H2 groups, respectively) were evaluated by TUNEL and CD8+ T cell staining. 4H2 was associated with an increase in tumor TUNEL signal and CD8+ T cell content by factors of 4.5±0.6 and 1.5±0.2, respectively, as compared to tumors in mice treated with IgG control (P<0.05) (**Fig. 7b, c, Supp. Fig. 2**). These findings suggest an immune mediated component of response to 4H2 in the PPQ model.

**Figure 7.**
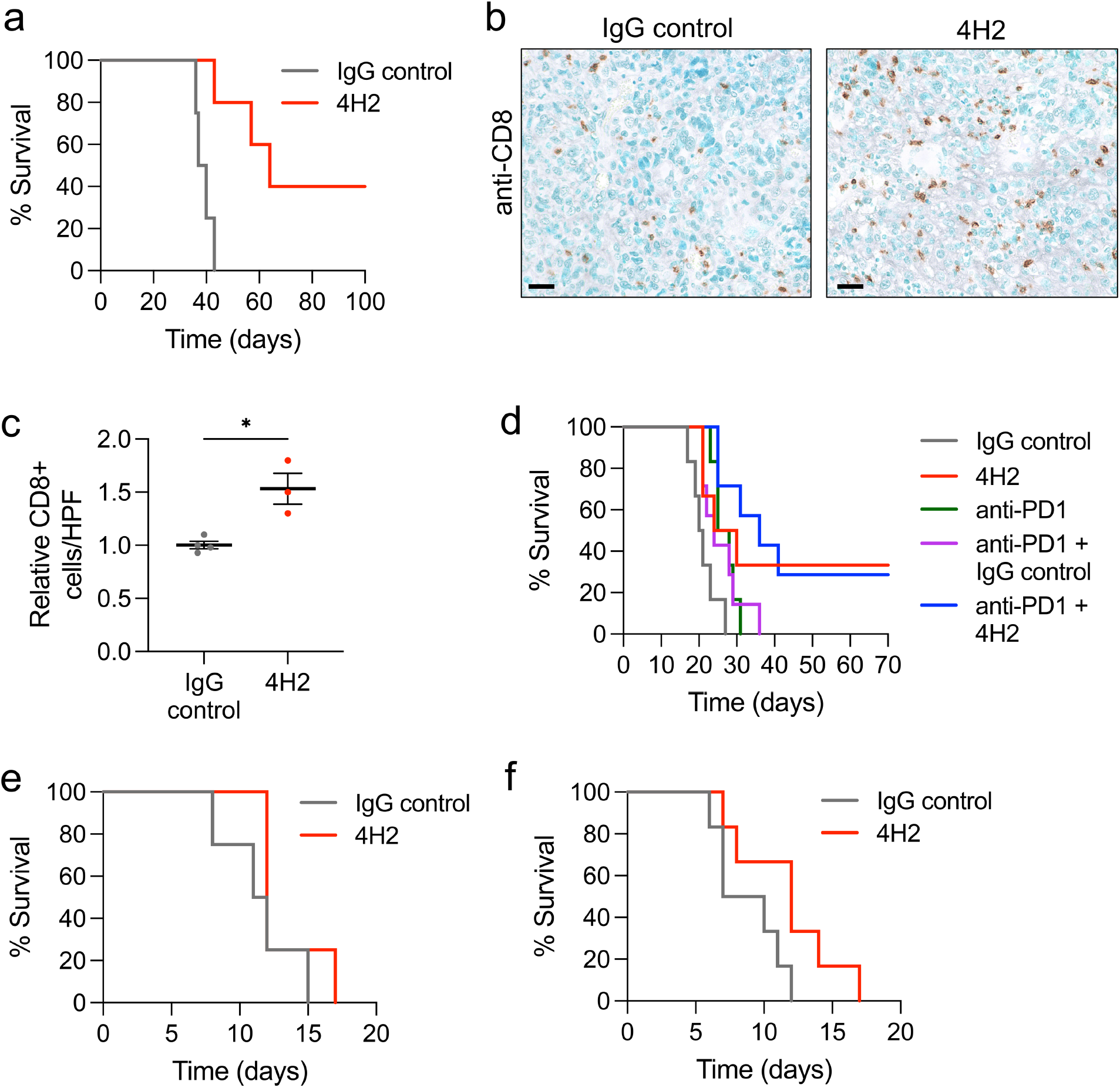
4H2 improves survival in syngeneic orthotopic GBM models. (**a**) Kaplan-Meier survival plots of C57/BL6 mice with PPQ orthotopic GBM brain tumors treated with three weekly cycles of IgG control (N=4) or 4H2 (N=5) are shown. 4H2 increased median survival by 66% compared to mice treated with IgG control (**P<0.01, log-rank test), and survival to study completion was 40% and 0% in groups treated with 4H2 or IgG control, respectively. (**b, c**) 4H2 increases CD8+ T cell infiltration into tumors. Brain tumors from the mice described in (**a**) meeting criteria for euthanasia (N=4 and 3 for IgG control and 4H2 groups, respectively) were immunostained for CD8+ T-cells and relative CD8 counts/HPF were examined. Representative images are shown in (**b**) and counts in (**c**). 4H2 caused a relative increase in CD8+ T cell content in tumors by a factor of 1.5±0.2 compared to IgG control (*P<0.05). (**d**) Kaplan-Meier survival plots of mice with GL261 orthotopic GBM brain tumors treated with IgG control (N=6), 4H2 (N=6), anti-PD1 (N=6), anti-PD1 + IgG control (N=7), or anti-PD1 + 4H2 (N=7). 4H2 increased median survival 32% compared to IgG control (*P=0.03, log-rank test) and when combined with anti-PD1 increased median survival by 50% compared to anti-PD1 + IgG control (*P=0.02, log-rank test). 4H2 alone and 4H2 + anti-PD1 yielded 33% and 29% survival to study completion, respectively, compared to 0% in all other groups. (**e, f**) 4H2 does not improve survival in an immunodeficient orthotopic GBM model. Kaplan-Meier survival plots of athymic nude mice with PPQ orthotopic GBM brain tumors treated with once (**e**) or twice weekly (**f**) cycles of IgG control (N=4 and 6, respectively) or 4H2 (N=4 and 6, respectively) are shown. 4H2 did not significantly impact median survival compared to IgG control in this immunodeficient model, demonstrating the importance of a functional immune system to the 4H2 effect on survival.

The GL261 syngeneic model is commonly used in pre-clinical studies of glioma immunotherapy and exhibits resistance to immune checkpoint blockade (ICB) (25-27). We selected GL261 as a second model for testing 4H2. Based on the enhanced immune response mediated by 4H2 in the PPQ model, we hypothesized that 4H2 would overcome GL261 resistance to immunotherapy and improve survival when administered alone or in combination with anti-PD1. C57/BL6 mice with GL261 orthotopic GBM tumors confirmed by IVIS were randomized to weekly treatment for three weeks with IgG control (N=6), 4H2 (N=6), anti-PD1 (N=6), anti-PD1 + IgG control (N=7), and anti-PD1 + 4H2 (N=7) and were monitored for toxicity and survival. No adverse effects were observed. 4H2 alone yielded a 32% increase in median survival compared to IgG control (P=0.03, log-rank test). When combined with anti-PD1, 4H2 increased median survival by 50% compared to anti-PD1 + IgG control (P=0.02, log-rank test). Groups treated with 4H2 alone or 4H2 + anti-PD1 showed 33% and 29% survival to study completion, respectively, compared to 0% in all other groups (**Fig. 7d, Supp. Table 2**). These findings demonstrate efficacy of 4H2 as a single agent and in combination with anti-PD1 against GL261 GBM tumors.

To test the importance of the immune system in mediating the anti-tumor effect of 4H2, athymic nude mice bearing intracranial PPQ GBM tumors were randomized to treatment once or twice weekly with IgG control (N=4 and 6, respectively) or 4H2 (N=4 and 6, respectively) and survival measured. No adverse effects of 4H2 were observed, and 4H2 did not yield any significant improvement in median survival compared to IgG control. Survival to study completion was 0% in all groups (**Fig. 7e, f, Supp. Table 3, 4**). These results indicate that the anti-tumor effect of 4H2 *in vivo* is dependent on the presence of a functional immune system.

## Discussion

This study reveals a previously unknown interaction between a lupus anti-DNA autoantibody and the cytoplasmic DNA sensor cGAS. Specifically, anti-DNA autoantibody 4H2 penetrates cells and localizes to cytoplasm through a GUO and nucleoside transporter-dependent mechanism, avoids endosomes, activates cGAS, causes cGAS-dependent toxicity to glioma cells, and promotes an immune-mediated prolongation of survival in orthotopic GBM models. These findings open new opportunities to apply cytoplasmic-localizing anti-DNA autoantibodies in cancer immunotherapy.

The BBB presents a significant challenge in systemic approaches to brain tumor therapy. We recently discovered a novel nucleoside transporter-dependent method of BBB crossing that facilitates brain tumor-targeting by systemically administered anti-DNA autoantibody DX1 (5). The present work now extends this mechanism to 4H2. 4H2 transport into cells and across a transwell model of the BBB was linked to nucleoside transport, evidenced by the respective enhancing and inhibitory effects of GUO and the nucleoside transport inhibitor DP. Further, systemically administered 4H2 localized to brain tumors where DNA/nucleosides are released, but not to normal brain tissue. The preferential tumor penetration by 4H2, with sparing of normal brain and otherwise minimal deposition in kidneys over IgG control, mitigates concerns for off-target toxicity and parallels the tumor-targeting behavior exhibited by DX1 (5). cGAS-STING agonists have proven effective against GBM in pre-clinical work, but direct intracranial and/or nanoparticle-based delivery of drug was required (28-30). The ability of systemically administered 4H2 to localize to brain tumor and activate cGAS suggests its potential for use in brain tumor immunotherapy.

Theories on mechanisms of autoimmunity are continually evolving, and here we introduce the new concept of cGAS activation by a cytoplasmic-localizing anti-DNA autoantibody. In addition to the translational applications in cancer immunotherapy discussed above, this finding raises the possibility that cGAS-mediated immune activation by cell-penetrating autoantibodies may contribute to mechanisms underlying inflammation and the interferon signature in SLE (2, 31). Moreover, this effect may play a role in protecting against some malignancies in SLE through enhancing tumor immunity (1, 2). Nuclear-penetrating anti-DNA autoantibodies were previously shown to cause DNA damage in cancer cells (2-5), and in this work a cytoplasmic-localizing variant is now found to activate cGAS. Overall, the diverse epitope binding and effects on cancer cells and the immune system associated with cell-penetrating anti-DNA autoantibodies make them compelling biologics for use in cancer immunotherapy.

## Materials and Methods

### Hybridomas and cell lines

4H2 hybridoma was obtained under MTA with the University of California, Los Angeles. 4H2 IgG2a antibody was purified from hybridoma supernatant as previously described (18). For control, mouse IgG2a monoclonal antibody (IgG control) was purchased from Leinco Technologies (I-118). DX1 was purified as previously described (5). Mouse anti-PD1 antibody was purchased from Bio X Cell. Cal12T cells were obtained from Horizon Discovery Ltd (Cambridge, UK), and U87 and A549 cells from the ATCC. hCMEC/D3 cells were purchased MilliporeSigma (SCC066) and NHA from Lonza as previously described (12). PPQ cells were obtained from MD Anderson Cancer Center (21). Cells were grown in 10% FBS supplemented RPMI 1640 and maintained at 37°C/5% CO_^2^_.

### Western blotting

Cells were treated with media containing specified amounts of control media, IgG control, or 4H2 overnight, after which cell lysates were prepared, subjected to 4-15% SDS-PAGE, and nitrocellulose transfer. Membranes blocked with 5% milk in TBST were incubated with relevant primary antibodies overnight at 4°C, washed and incubated with HRP-conjugated anti-rabbit or mouse IgG secondary antibodies (Cell Signaling) for one hour at room temperature. After additional washing, bands were detected by Lumiglo (Cell Signaling Technologies).

### Cell penetration and co-localization immunofluorescence assays

Live cells were grown in 96-well plates or on glass coverslips and treated with control media or media containing 0.5-1 mg/mL IgG control, DX1, or 4H2 for one hour. Cells were then washed and fixed, and intracellular location of 4H2 or IgG control was detected by immunostaining with an Alexa Fluor 488-conjugated goat anti-mouse IgG antibody (Cell Signaling, Danvers, MA) as previously described (32), and DX1 immunostaining as previously described (5). Western blots on Cal12T lysates were probed with HRP-linked horse anti-mouse IgG (Cell Signaling, #7076) and mouse anti-β-actin (Ambion, Austin, TX). For co-localization studies, the fixed cells were also probed overnight with rabbit primary antibodies to detect endosomes (C45B10 anti-EEA1 antibody, Cell Signaling), lysosomes (D2D11 anti-LAMP1 antibody, Cell Signaling), endoplasmic reticulum (C81H6 anti-PDI antibody, Cell Signaling), Golgi apparatus (D2B6N anti-RCAS1 antibody, Cell Signaling), mitochondria (3E11 anti-COX IV, Cell Signaling), after which they were washed with PBS and incubated with an Alexa Fluor 555-conjugated goat anti-rabbit IgG antibody (Cell Signaling) at room temperature for one hour. After a final series of washes, cells were treated with Prolong Gold Antifade Reagent with DAPI (Cell Signaling) and imaged using an EVOS fl digital fluorescence microscope (Advanced Microscopy Group, Bothell, WA) under light, DAPI, GFP, or RFP filters. Fluorescence images were merged using ImageJ (NIH, Bethesda, MD). For DP studies, cells were pre-treated with control media or media containing 50 µM DP (MilliporeSigma, D9766) for thirty minutes, followed by addition of 1 mg/mL 4H2 for one hour, and then evaluated for cellular penetration as described above. 4H2 fluorescence intensity was quantified using ImageJ.

### FITC-labeling of 4H2 and live cell imaging

Purified 4H2 was labeled with FITC using the Pierce FITC Antibody Labeling Kit (Thermo Fisher Scientific, Waltham, MA). For the live cell penetration assays Cal12T and A549 cells were treated with FITC-labeled 4H2 at 0.5 mg/mL overnight, and then treated with 500 nM MitoTracker Red FM (Thermo Fisher Scientific) for 45 minutes followed by 1 µg/mL Hoechst 33342 (Thermo Fisher Scientific) for 15 minutes. Cells were then washed with PBS and imaged using the EVOS fl digital fluorescence microscope.

### Purified cGAS binding studies

2 µg recombinant human cGAS (Cayman Chemical, #22810) was incubated with 8 µg IgG control or 4H2 ± nucleic acid (1.25 µg PvuII-digested pcDNA3 and 50 µM GTP) in a volume of 500 µL PBS with 0.01% Triton X-100 for one hour at 4°C with rotation. 50 µL of 50% protein G bead slurry was added to the reaction and incubation continued for one hour at 4°C with rotation. Supernatant was removed and beads were washed five times in PBS + 0.05% Triton X100, and bound protein eluted in 50 µL Pierce Gentle Ab/Ag Elution Buffer (Thermo Fisher Scientific #21027). Samples were analyzed by western blot probed with rabbit monoclonal antibody against human cGAS (Cell Signaling, #15102) and HRP-linked goat anti-rabbit IgG (Abcam, ab205718).

### Cell lysate antibody pulldown assays

PPQ cells treated with 1 mg/mL IgG control or 4H2 for one hour were washed and contents harvested using NE-PER™ Nuclear and Cytoplasmic Extraction Kit (Thermo Fisher, #78833). 50 µL 50% protein G bead slurry was added to the 500 µL final extraction volume for incubation at 4°C with rotation for 1.5 hours. Beads were washed, and remaining bound proteins eluted in 40 µl SDS loading sample buffer. Samples were analyzed by western blot probed for cGAS with rabbit anti-mouse cGAS (Cell Signaling #31659) at 1:1000 and secondary goat anti-rabbit HRP (abcam ab205718) at 1:5000.

### cGAS activity assay

The effect of IgG control or 4H2 (0-160 µg) on cGAMP production was assayed using the cGAS Inhibitor Screening Assay Kit (Cayman Chemical, #701930) as per the manufacturer’s instructions except we used half of the recommended quantity of cGAS to allow room to detect signal amplification in the presence of 4H2. The ratio of cGAMP produced in the presence of 4H2 to cGAMP made in the presence of IgG control was determined.

### NF-κB assay

PPQ cells were treated with 1 mg/ml IgG control or 4H2 for 18 hours in growth medium (DMEM+10% FBS). Cytoplasmic and nuclear fractions were harvested using with NE-PER™ Nuclear and Cytoplasmic Extraction Kit (Thermo Fisher Scientific, #78833) and evaluated by NF-κB (Cell Signaling, #8242) western blot with Lamin B1 (Cell Signaling, #12586) for loading control. Band intensities were determined using ImageJ.

### cGAS knockdown and colony formation assays

PPQ cells grown in 6-well plates were transfected with 100 nM control or cGAS siRNA (Dharmacon) by RNAiMax (Thermo Fisher Scientific). cGAS knockdown was confirmed by western blot three days later. On the same day cells were treated with IgG control or 4H2 (0-1.6 µM) and evaluated for clonogenic survival by colony formation assay.

### Nucleoside supplementation assays

PPQ cells grown in 96-well plates were pre-incubated for thirty minutes in cell culture media ± 100 µM ADE or GUO. Cells were then treated by addition of control media or media containing DX1 ± 100 µM ADE or 4H2 ± 100 µM ADE or GUO for five minutes, after which media was withdrawn, cells were washed, fixed, and immunostained to detect DX1 or 4H2. Cell staining intensity was then quantified using ImageJ, and relative uptake in the presence or absence of nucleoside was determined.

### Transwell model of the BBB

The ability of 4H2 to cross a transwell model of the BBB was tested using our previously described protocol (5). hCMEC/D3 brain endothelial cells and NHA were respectively adhered to apical and basolateral sides of cell culture inserts (MilliporeSigma 353095) coated with fibronectin (MilliporeSigma F1141) and poly-L-lysine (MilliporeSigma P4832). Formation of a functional barrier was confirmed as previously described (5), and BBB models treated with control buffer or 50 µM DP for 30 minutes followed by 5 µM 4H2 ± 50 µM DP. Relative 4H2 crossing of the barrier at 15 and 30 minutes in the presence or absence of DP was evaluated by ImageJ quantification of anti-mouse IgG dot blots on basolateral chamber contents.

### Orthotopic GBM studies

Studies were conducted under a Yale University IACUC approved protocol. Orthotopic GBM tumors were established in female C57/BL6 mice aged 5-6 weeks by stereotactic injection of 50,000 cells (PPQ or GL261 engineered to express luciferase) and tumor formation verified by IVIS Spectrum *in vivo* imaging system (IVIS) using our previously described method (5). Mice with tumors confirmed by IVIS were randomized to treatment with tail vein injection of IgG control (40 mg/kg tail vein) or 4H2 (40 mg/kg tail vein) once per week for three weeks in the PPQ study, and IgG control (40 mg/kg tail vein), 4H2 (40 mg/kg tail vein), anti-PD1 (5 mg/kg IP), and combinations thereof as described in the results in the GL261 study. Mice were closely monitored throughout the study and were humanely euthanized for endpoints of neurologic change or weight loss. Kaplan Meier survival plots and median survivals were generated using GraphPad Prism version 9.4.1. For antibody localization studies, tumors and normal tissues were harvested 24 hours after a single tail vein injection of IgG control or 4H2 (40 mg/kg), fixed in 10% neutral buffered formalin, and paraffin embedded. Presence of antibody was examined by IHC probed with anti-mouse IgG-HRP at 1:50 using our previously described protocols (5). CD8+ T cell content in tumors was evaluated using anti-CD8a at 1:2000 (abcam, ab217344) antibody and number of positive cells per HPF counted. Signal was developed with DAB and methylgreen counterstain. TUNEL stain was performed using Cell Signaling TUNEL Assay Kit (#25879) and signal quantified by ImageJ.

### Statistical analysis

Graphs were generated using GraphPad Prism version 9.4.1. P values were determined by one-tailed Student’s t-test or log-rank test where specified for *in vivo* studies. Error bars represent SEM.

## Acknowledgements

Supported by a Yale Cancer Center Pilot Research Grant, American Cancer Society Institutional Research Grant (#58-012-55), the Dept. of Therapeutic Radiology at Yale School of Medicine, and the National Institute of Neurological Diseases and Stroke of the NIH under award number R01NS112223 (JEH). The content is solely the responsibility of the authors and does not necessarily represent the official views of the NIH. Please note this manuscript is a preprint draft and has not yet completed peer review.

## Materials & Correspondence

Correspondence and requests for materials should be addressed to James E. Hansen.

