## Supplemental Figures and Tables for "cGAS-activating lupus autoantibody for cancer immunotherapy"

**Supplemental Figure 1**

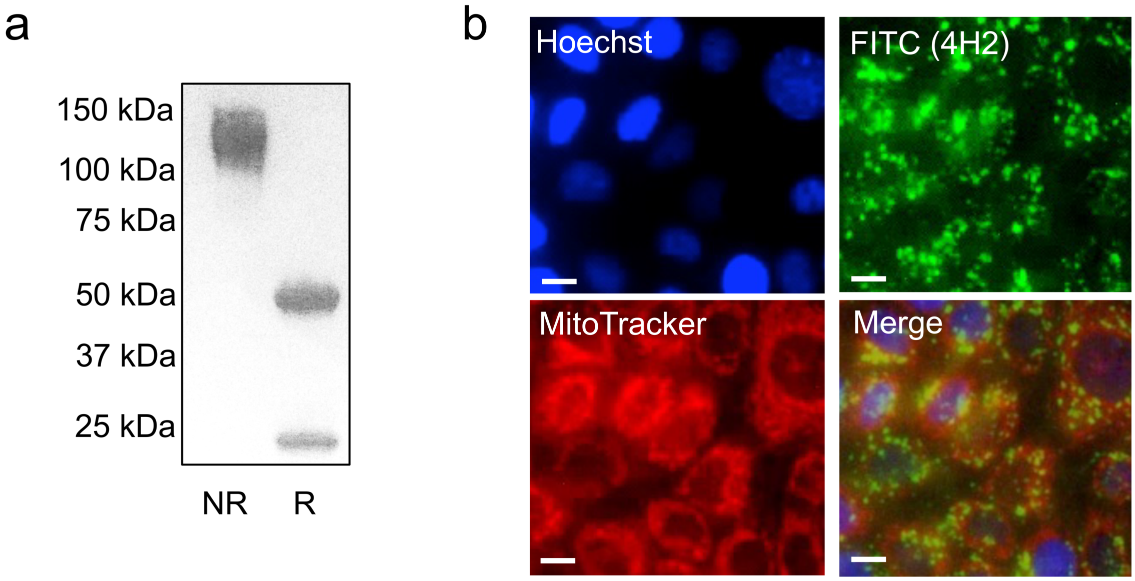


**Supplemental Figure 1. 4H2 penetrates live cells.** (**a**) Successful purification of 4H2 from hybridoma supernatant was confirmed by SDS-PAGE under reducing (R) and non-reducing (NR) conditions. Intact antibody and HC and LC were detected at their expected MWs. (**b**) Cal12T human lung cancer cells were treated with 0.5 mg/mL FITC-labeled 4H2 overnight, and cells were then co-stained with MitoTracker Red FM for detection of mitochondria and with Hoechst for nuclear visualization. Live cells were visualized under a fluorescence microscope, and representative images are shown. 4H2-FITC signal was detected in the cytoplasm. Bars =10 μm.

**Supplemental Figure 2**

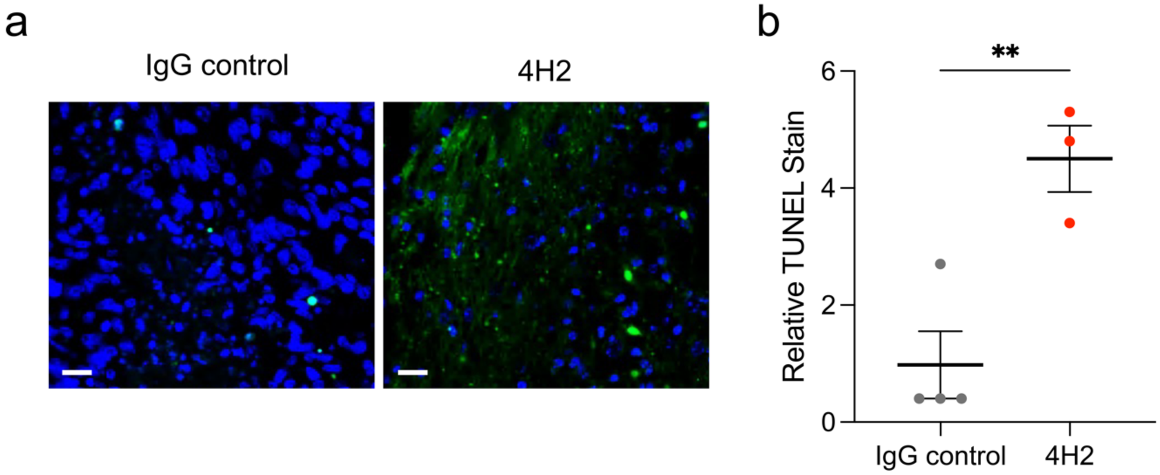


**Supplemental Figure 2. TUNEL staining is increased in GBM tumors in mice treated with 4H2.** (**a**) Sections of PPQ GBM tumors in C57/BL6 mice treated with IgG control or 4H2 were analyzed by TUNEL (green) and DAPI (blue) staining. Representative merged fluorescence images are shown. Adjustments to brightness/contrast were applied equally to IgG control and 4H2 image sets. Bar: 20 μm. (**b**) Quantification of TUNEL staining by ImageJ showed a relative fold increase in TUNEL signal of 4.5±0.6 in mice treated with 4H2 compared to IgG control (**P<0.01).

**Supplemental Table 1**

| Group | Median Survival | % Survival at 100 days |
|---|---|---|
| IgG control | 38.5 days | 0 |
| 4H2 | 64 days | 40 |

**Supplemental Table 1.** Median and percentage survival at study completion in C57/BL6 mice with PPQ orthotopic GBM tumors treated with IgG control or 4H2. 4H2 increased survival compared to IgG control ($P<0.01$, log-rank test).

**Supplemental Table 2**

| Group | Median Survival | % Survival at 70 days |
|---|---|---|
| IgG control | 20.5 days | 0 |
| 4H2 | 27 days | 33 |
| anti-PD1 | 26.5 days | 0 |
| anti-PD1 + IgG control | 24 days | 0 |
| anti-PD1 + 4H2 | 36 days | 29 |

**Supplemental Table 2.** Median and percentage survival at study completion in C57/BL6 mice with GL261 orthotopic GBM tumors treated with combinations of IgG control, 4H2, or anti-PD1. 4H2 increased survival compared to IgG control ($P=0.03$, log-rank test). Addition of 4H2 to anti-PD1 increased survival compared to anti-PD1 alone ($P=0.02$, log-rank test) or to anti-PD1 + IgG control ($P=0.02$, log-rank test).

**Supplemental Table 3**

| Group | Median Survival | % Survival at 20 days |
|---|---|---|
| IgG control | 11.5 days | 0 |
| 4H2 | 12 days | 0 |

**Supplemental Table 3.** Median and percentage survival at study completion in athymic nude mice with PPQ orthotopic GBM tumors treated with weekly IgG control or 4H2. 4H2 did not change survival compared to IgG control (P=ns, log-rank test).

**Supplemental Table 4**

| Group | Median Survival | % Survival at 20 days |
|---|---|---|
| IgG control | 8.5 days | 0 |
| 4H2 | 12 days | 0 |

**Supplemental Table 4.** Median and percentage survival at study completion in athymic nude mice with PPQ orthotopic GBM tumors treated with twice weekly IgG control or 4H2. 4H2 did not change survival compared to IgG control (P=ns, log-rank test).
